# Information Integration and Collective Motility in Phototactic Cyanobacteria

**DOI:** 10.1101/590778

**Authors:** S. N. Menon, P. Varuni, G I. Menon

**Affiliations:** The Institute of Mathematical Sciences, C.I.T Campus, Taramani, Chennai 600113, Tamil Nadu, India

**Author notes:** These authors contributed equally to this work (S. N. Menon, P. Varuni).

## Abstract

Cells in microbial colonies integrate information across multiple spatial and temporal scales while sensing environmental cues. A number of photosynthetic cyanobacteria respond in a directional manner to incident light, resulting in the phototaxis of individual cells. Colonies of such bacteria exhibit large-scale changes in morphology, arising from cell-cell interactions, during phototaxis. These interactions occur through type IV pili-mediated physical contacts between cells, as well as through the secretion of complex polysaccharides (‘slime’) that facilitates cell motion. Here, we describe a computational model for such collective behaviour in colonies of the cyanobacterium *Synechocystis*. The model is designed to replicate observations from recent experiments on the emergent response of the colonies to varied light regimes. It predicts the complex colony morphologies that arise as a result. We ask if changes in colony morphology during phototaxis can be used to infer if cells integrate information from multiple light sources simultaneously, or respond to these light sources separately at each instant of time. We find that these two scenarios cannot be distinguished from the shapes of colonies alone. However, we show that tracking the trajectories of individual cyanobacteria provides a way of determining their mode of response. Our model allows us to address the emergent nature of this class of collective bacterial motion, linking individual cell response to the dynamics of colony shape.

**Statement of Significance:** Microbial colonies in the wild often consist of large groups of heterogeneous cells that coordinate and integrate information across multiple spatio-temporal scales. We describe a computational model for one such collective behaviour, phototaxis, in colonies of the cyanobacterium *Synechocystis* that move in response to light. The model replicates experimental observations the response of cyanobacterial colonies to varied light regimes, and predicts the complex colony morphologies that arise as a result. The results suggest that tracking the trajectories of individual cyanobacteria may provide a way of determining their mode of information integration. Our model allows us to address the emergent nature of this class of collective bacterial motion, linking individual cell response to the large scale dynamics of the colony.

## Introduction

Cells respond to a variety of sensory inputs, including chemical and physical signals. An experimentally measurable example of such behaviour involves cell motility, where cells alter their motion in response to an external signal [1]. Bacteria provide a particularly convenient model to investigate taxis to many types of stimuli, including pH changes [2], oxygen [3], osmolarity [4] and magnetic fields [5]. Chemotaxis, where cells swim up (or down) chemical gradients, is an extensively studied example of cell taxis, most notably in flagellated *Escherichia coli*. However, while the responses of individual cells to single inputs have been well characterized and modeled [6], the mechanisms through which cells collectively respond to more complex and spatially structured combinations of inputs remain open to investigation.

Cyanobacteria exhibit phototaxis, or motion in response to a light stimulus [7]. When colonies of the model cyanobacterium *Synechocystis* sp. PCC 6803 are exposed to red or green light emanating from a single source, individual cells first move toward the edge of the colony nearest to the light source. There, they aggregate before further extending towards the source through regular, dense finger-like projections [8]. Variations in light intensity and wavelength induce responses that range from slower moving colony fronts [8] to negative phototaxis [9]. Phototactic cells such as *Synechocystis* respond directly to the relative position of the light source [10] and not to a spatio-temporal concentration gradient, as in the case of chemotaxis.

Unlike flagellar-driven motion of *E. coli*, *Synechocystis* exhibits “twitching” or “gliding” motility is slower and has lower directional persistence [11]. This mode of motility is facilitated by type IV pili (T4P). These pili attach to the substrate and retract to move the cell forward [11]. This type of motility is often also associated with complex polysaccharides, or ‘slime’, extruded by these cells. The presence of slime reduces the friction that cells experience during motion [12]. The T4P also add another collective component to gliding motility, since cells can also use them to attach to each other [11]. Further, while *E. coli* provides an example of a single-cell response that can be studied at high resolution, cells in their natural environments are often found in dense aggregates and biofilms where interactions between cells are harder to probe, yet cannot be ignored.

The non-linear collective response arising from cell-cell communication, as in quorum sensing, provides an example of how interactions between cells drives qualitatively different behaviour [13]. These types of collective behaviour are often hard to capture in single cell models. Further, both light quality and direction can fluctuate in the natural environment, but the effects of such variation are not currently well understood. Several studies have explored the effects of varied illumination schemes on colony morphology [14–19]. In one recent experiment [19], colonies of *Synechocystis* receive light incident on them from two different directions. These studies found that fingers from the colonies emerged along a direction intermediate between the directions of the light sources.

This raises the question of whether this collective behaviour at the level of the entire colony is best interpreted as arising from individual cells attempting to move along an intermediate direction determined by the vector sum, or whether it could also plausibly arise from the averaged response of individual cells responding to a single, randomly chosen light source at each time. Identifying which of these two scenarios occurs would require experiments that probe the coupled dynamics of receptor activation and downstream signaling, with consequences for cell motility. Such relationships are hard to establish in practice, and it is thus worthwhile to explore alternative ways of discriminating between such scenarios, guided by modeling.

Here, we quantitatively analyse the different modes of individual cell behaviour and their resultant colony morphologies in the context of a model for the light-directed motion of cells in cyanobacterial colonies. Our perspective on such collective phenomena is motivated by models for active matter [20, 21], which describe the collective behaviour of systems of self-propelled units. Our earlier model [22] incorporated motion at the level of single cells, cell-cell interactions mediated via T4P and the decrease in surface friction through the deposition of slime. That model is extended here to incorporate what is known about the response of cells to variations in light intensity and wavelength, including the possibility of negative phototaxis.

Our modeling framework addresses recent experiments involving the complex illumination of cyanobacterial colonies. We reproduce experimental results and demonstrate how our model can be generalized to novel situations involving several light sources, each with a different wavelength and intensity. The model makes predictions regarding the trajectories of individual bacteria in cyanobacterial colonies. Importantly, we find that colony morphology cannot be used to uniquely infer mechanisms through which individual cells integrate information from multiple light sources, since the large-scale morphology of the colony is independent of whether individual cells decide to move along the vector sum of the light they receive, or whether they make stochastic decisions to move towards one or the other of these inputs at each step. However, examining the trajectories of single cells within such colonies provides a way of distinguishing between different scenarios for information integration. Furthermore, we find that qualitatively similar results are obtained even when individual cells respond heterogeneously. This is largely a consequence of the fact that the motion of groups of cells that interact with each other involves a collective component.

## Methods

The model used in this paper is adapted from a previously proposed model [22] for the collective motion of cyanobacterial colonies illuminated by a single light source. This model described the behaviour of independently motile cells that can physically interact with each other, and move towards a distant light source. The three essential components of this model are: (i) the ability of cells to locate the position of a light source that biases their direction of motion; (ii) the forces that cells exert on other cells in their vicinity through T4P, and; (iii) the deposition of slime by individual cells, which reduces the friction that they encounter.

In this paper, we extend this earlier model to account for complex illumination arising from multiple light sources. Our current model describes two scenarios for how cells might integrate directional input in the form of separated light sources to which they respond. We use LED as a representation for a generic light source in all our figures. The schematic of Fig. 1(a) describes the general setup of our simulations involving cell colonies subjected to two light sources, whose individual intensities and wavelengths may vary. We briefly describe this model below.

**Figure 1:**
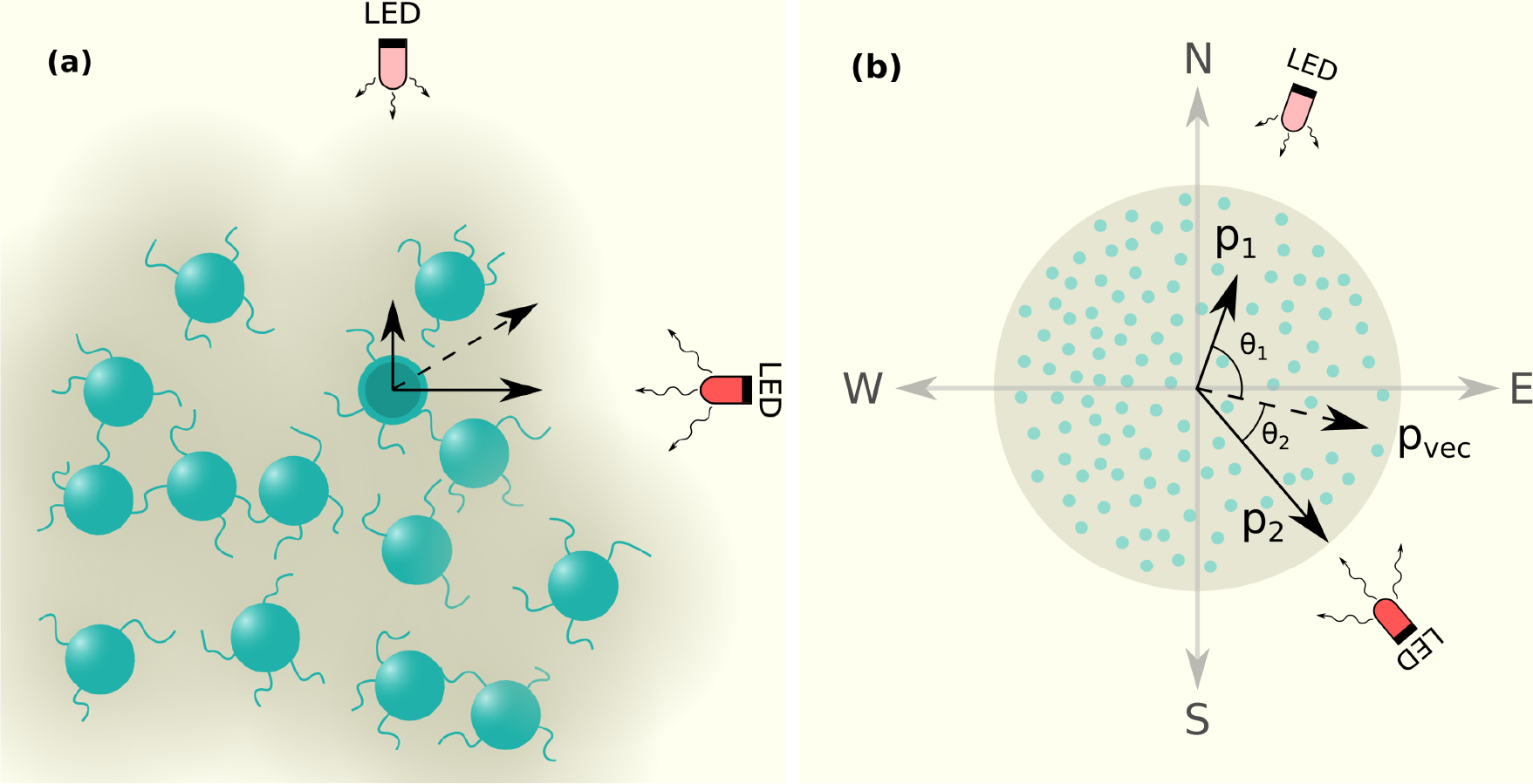
Schematic of simulation system. (a) Cells attached to the substrate are shown as green circles. The intensity of the background color (gray) represents the amount of underlying slime. Cells can attach to other cells through T4P, shown as thread like extensions from the cells. The two light sources (LED) are positioned as shown. (b) A cell colony containing multiple cells and exposed to two light sources of different intensities, specified via *p*_1_ and *p*_2_, as shown. Directions are specified as (N, S, E, W). The angles made by the sources with the E-W axis (horizontal) are indicated as *θ*_1_ and *θ*_2_. The quantities *p*_1_ and *p*_2_ represent probabilities of moving in the direction of either source independently, so can be thought of as vectors with the magnitude *p*_1_ and *p*_2_ aligned with the radius vector from the colony centre to the light source. The resultant *p*_vec_ represents the vector sum of these quantities. In all subsequent figures that display the morphologies of colonies obtained from simulations, we use a similar visual representation, i.e. individual cells are represented by green circles, and the amount of underlying slime is represented by the intensity of gray.

### Modeling complex illumination

We consider arrangements of individual colonies that receive light from sources placed at different locations. The light from each source may also present a different wavelength. The intensity arising from each light source is assumed to be uniform within a colony but can vary across colonies. The angle that the line joining the colony centre to a light source *k* makes with the horizontal is given by Θ_*k*_, as illustrated in the schematic of Fig. 1(b). For ease of description, we label the directions as (North, South, East, West), or (N, S, E, W) as shown in Fig. 1(b). In all cases, each light source is placed one unit away from the grid that defines the colony locations. The probability that cells within a given colony attempt to move towards light source *k*, in the Θ_*k*_ direction is captured by *p*_*k*_. The schematic of Fig. 1(b) shows two light sources separated by an arbitrary angle, leading to an inhomogeneous distribution of intensities across the colony.

We replicate the experimental setup of [19] as closely as possible, as illustrated in Fig. 1(b). In these experiments, the intensity of light experienced by a colony varies with its distance from the source. We assume that *p*_*k*_ for each light source varies inversely with the distance, *d*_*k*_, between the center of the colony and the light source.

### The cell

Each cell is modeled as a disc of radius R, specified by a two dimensional vector, *X*_i_ = (*x*_i_, *y*_i_). As in [22], when colonies are subjected to a single light source *k*, individual cells attempt to move in the direction Θ_*k*_ with probability *p*_*k*_ or in a random direction in the interval [0, 2*π*] with probability 1 − *p*_*k*_.

We extend this model to include illumination from multiple light sources. Fig. 2 displays two possible ways in which individual cell movement in such colonies may be biased towards two distinct light sources, placed to the North and to the East of a colony. The first column (Fig. 2(a) and Fig. 2(b)) illustrates the case where the sources are switched on individually, while the second column (Fig. 2(c) and Fig. 2(d)) illustrates two distinct cell responses in the case where both light sources are switched on, viz. stochastic switching and vector integration.

**Figure 2:**
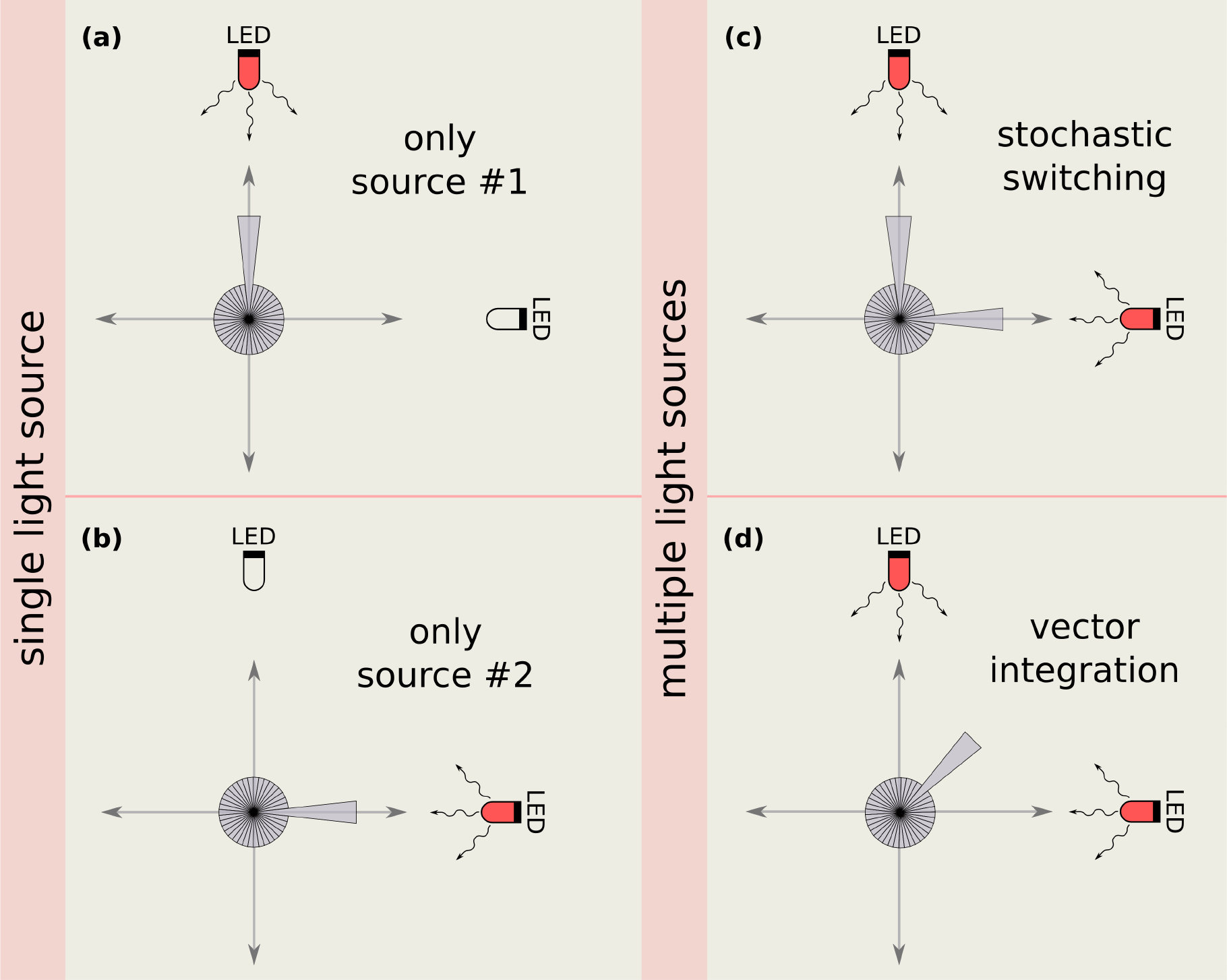
Possible biases of individual cells under a complex illumination scheme. In the presence of a single light source, cells sense the light and their direction of motion is biased towards it, as in (a) and (b). The length of each wedge represents the probability that a cell moves along the direction of the centre line of the wedge. When no external source of light is present, all wedges have the same size. In the presence of an external light source, the wedge in the radial direction of that source is enlarged, since the probability of moving in that direction is increased. When cells are subjected to two different light sources, as shown in panels (c) and (d), there are two possible mechanisms for them to integrate this information. Either, as shown in the schematic (c) they can stochastically choose, at every step, to bias their motion towards one of the two light sources or, as shown in (d), they can bias their motion towards the vector sum of the directions of the two light sources.

#### Stochastic switching

At each time step, cells may decide to move in the direction Θ_*k*_ of any of the light sources *k* with probability *p*_*k*_, or in a random direction in the interval [0, 2*π*] with probability 1 − ∑*p*_*k*_. This is illustrated in Fig. 2(c).

#### Vector integration

The vector joining the centre of each colony at *t* = 0 with light source *k* is represented by *v_k_* = *p_k_*(cos Θ_*k*_, sin Θ_*k*_). Individual cells attempt to move along the vector sum of these light sources, *v*_vec_ = *p*_vec_(cos Θ_vec_, sin Θ_vec_) = ∑ *v*_*k*_. Thus, cells attempt to move in the direction Θ_vec_ with probability *p*_vec_, or in a random direction in the interval [0, 2*π*] with probability 1 − *p*_vec_. This is illustrated in Fig. 2(d).

In both mechanisms, each cell *i*, at time *t*, picks a direction, 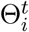 as described above. The decision of the cell to move in a particular direction is modeled through a force

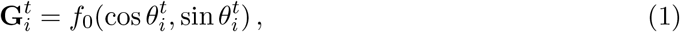

where we choose *f*_0_ = 1 to set the scale of forces. The direction in which a given cell actually moves is determined both by this chosen direction as well as by the forces it experiences from other cells in its neighbourhood. These forces, arising from the interaction between cells, are described below.

### Cell-cell interactions

Each cell is assumed to have a fixed number *m* of T4P. These can exert forces on randomly chosen cells that lie within a certain distance *ℓ* of its centre. During each time step, a cell *j* can exert a force *f*_*ji*_ = *K*_*ji*_(cos *θ*_*ji*_, sin *θ*_*ji*_) on a randomly chosen neighbouring cell *i* where *θ*_*ji*_ is the angle that the vector from cell *i* to *j* makes with the horizontal. The magnitude *K*_*ji*_ of this force depends on the distance *D*_*ji*_ between the cells *i* and *j*:

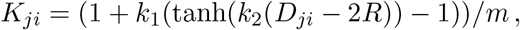

We use a sigmoidal form for *K*_*ji*_ that is repulsive at short distances. This penalizes cell overlaps through a soft-core repulsion. A detailed discussion of the motivation for choosing this specific form of *K*_*ij*_, and of the associated parameter values, is presented in [22].

The cell *i* thus experiences a total external force,

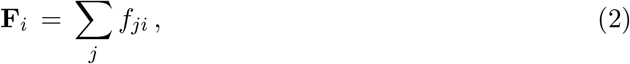

from other cells *j* in its neighbourhood. The net force acting on this cell at each time step *t* is then 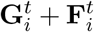.

### Slime Deposition

The slime deposited by cells is assumed to be deposited on a regular square lattice underlying the colony. Each grid point is specified by (*r, c*). Cells are assumed to deposit slime at every time step. The amount of slime, *S*^*t*^ at time *t* associated with the grid point closest to each cell’s centre, is incremented by an amount *S*_rate_ in each time step.

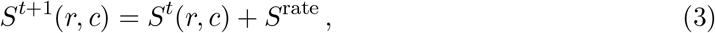

We further assume that once the amount of slime at a grid point exceeds *S*_max_, no more slime is added to that point. Additionally, we assume that slime does not decay or diffuse once deposited.

### Cell movement

The motility of a cell depends on the amount of slime at the grid point closest to the cell centre. The positions of cells are updated in parallel, as in standard agent-based models, with parameters chosen such that the maximum distance that a cell in a slime-rich background can move in a single time step is a tenth of the cell radius. In slime-poor backgrounds, the reduced mobility of the cell implies that it moves a smaller distance in the same time. At the end of each time step, the position of each cell is updated through the following scheme:

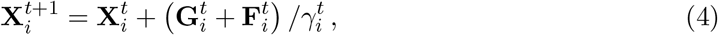

where 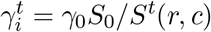 is a friction factor that is associated with the presence of slime lying below a cell, where *S*_0_ is the initial slime concentration within the colony and γ_0_ is the friction encountered by cells within the colony at *t* = 0.

### Simulation details

We simulate cyanobacterial colonies containing 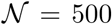 cells that are initially distributed randomly over a circular spatial domain representing a colony. We have verified that the results are qualitatively invariant for larger system sizes (see Supplementary Information Figs. S1 and S2 for results obtained for 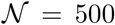, 2000 and 5000 over a range of values of *p*_photo_). In experiments on phototaxis the number of cells in a colony is typically of the order of tens of thousands of cells. Specifically in [19] each colony, which is around 2.5*mm* in diameter, consists of ~ 33000 cells, which is less than one order of magnitude higher than the maximum number of cells considered in our simulations. The parameter of significance here is the density *ρ* which (Supplementary Information Fig. S3), in experiments, is roughly 0.06 as calculated from the methods described in [19] and is similar to the *ρ* = 0.1 used in these simulations.

Unless otherwise indicated the parameters used in our simulations are listed in Table 1. *Synechocystis* cells are around 1*μm* [7] in radius, and we use this to define our cell radius *R*, which is also assumed to be the basic length scale in our model. Each cell in our model can have up to *m* appendages, and in these simulations, we use *m* = 4 [23]. The maximum length *l* of an appendage is taken to be four cell lengths [24]. For a detailed discussion of the cell force parameters (*k*_1_, *k*_2_), see [22]. Cells can move at a maximum of 0.1 body lengths per unit time [11]. To our knowledge there has not been a detailed investigation of slime deposition rate and how this affects cell speed. For a more detailed discussion of the system parameters see [22].

**Table 1:** Parameters used in simulations (unless mentioned otherwise)

| Parameter | Quantity | Value |
| --- | --- | --- |
| $\mathcal{N}$ | number of cells | 500 |
| $R$ | cell radius | 1 |
| $\rho$ | cell density in colony | 0.1 |
| $\ell$ | appendage length | $4 R$ |
| $m$ | number of appendages per cell | 4 |
| $\gamma_0$ | inverse of initial cell speed | $1/(0.1 R)$ |
| $(k_1, k_2)$ | force parameters | (1, 2) |
| $S_{\text{rate}}$ | slime deposition rate | 0.1 |
| $p_{\text{photo}}$ | phototaxis probability | $(0, 0.05]$ |

At the start of each simulation, cells are distributed uniformly over a circular colony, where the initial slime concentration is the same for all grid points within the colony. At each time step for each cell we determine 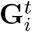 from Eq. (1). For each cell we also determine which of its neighbours it is attached to. Using this information, we can calculate the external force experienced by each cell 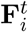 from Eq. (2). We then update the positions of each cell using the equation of motion (4), which involves the net force 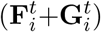 and slime underlying the cell. Finally, we update the slime matrix *S*^*t*^(*r, c*) using Eq. (3).

## Results

### Colony morphologies at varying light intensities

We consider colonies of cells that exhibit positive phototaxis. These are placed in a simple one-dimensional array consisting of 5 colonies, an arrangement similar to that used in the experiments of [19], and which is illuminated from the East. These experiments tested the response of colonies to varying intensities of red light. In our simulations, this variation in intensity is described by a single parameter, *p*_photo_, which varies across colonies.

Fig. 3 describes how the colony morphology changes as *p*_photo_ is varied between 0.05 and 0.01, simulating the drop in intensity as one moves from East to West across the arrangement of colonies. At smaller *p*_photo_, the colony emits small, slightly distorted fingers oriented towards the light source. The emerging fingers appear to meander more at low *p*_photo_ and the velocity of fingers in the direction of the source is reduced. As *p*_photo_ is increased, fingers become longer and more prominent. In addition, their velocity in the direction of the source is increased. The time-evolution of fingers at different *p*_photo_ values is illustrated in Supplementary Movie S1.

**Figure 3:**
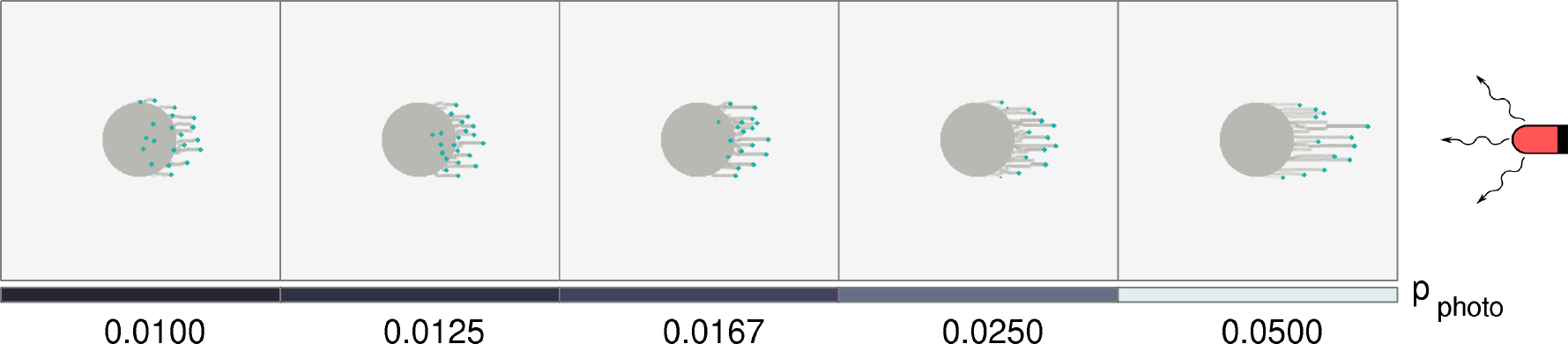
Cell colonies in a linear grid under complex illumination. Morphologies of several colonies under illumination from a single light source placed at the East. Each experiences a different intensity of incident light, depending on its location. The placement of the light source is such that the easternmost colony experiences the greatest incident light intensity. Thus, *p*_photo_ decreases linearly towards the West, as shown in the bottom bar. Each colony contains 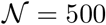 cells, and the model was simulated for 4 × 10^4^ time steps.

Our results, shown in Fig. 3, recapitulate the following experimental observations: (i) there is a light-flux dependent increase in the movement bias of cells in colonies, and (ii) finger sizes decrease at lower illumination.

### Colony morphologies under multiple light sources

We generalize the linear complex illumination described in the previous section by simulating intensity variation along two directions, an arrangement that was also considered in the experiments of [19]. In these experiments, colonies were arranged on a grid and exposed to two sources of red light, placed North and West of this grid.

In Fig. 4 and Fig. 5, we consider a set of 25 colonies placed in a 5 × 5 array and illuminated by two light sources. The intensities decrease along E-W and N-S directions as one moves away from each light source. The rules by which individual cells respond to this complex illumination can be either through Stochastic switching (Fig. 4) or Vector integration (Fig. 5) mechanisms, as described below.

**Figure 4:**
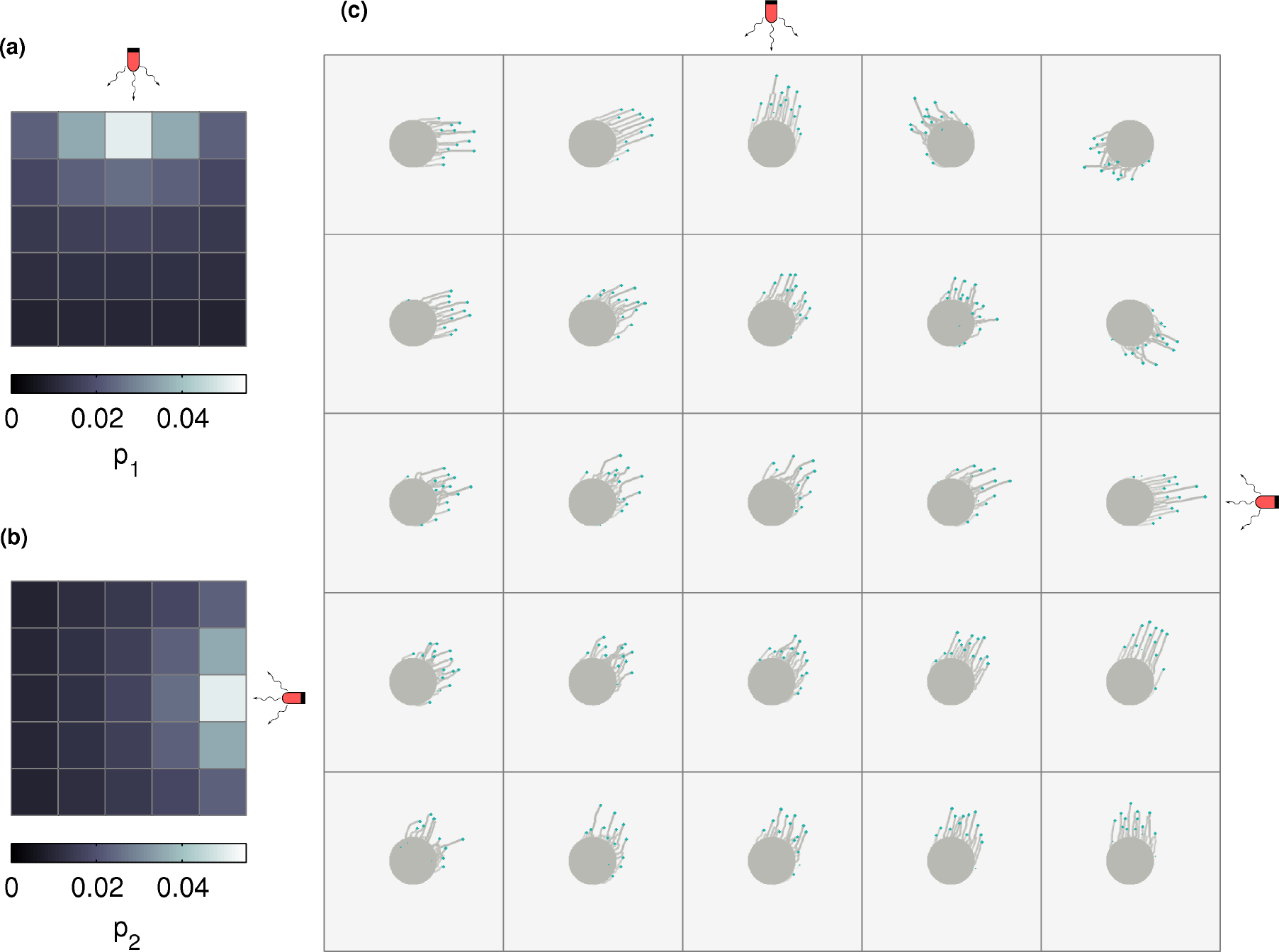
Cell colonies in a square grid under complex illumination in the stochastic switching case. Configurations of a number of cell colonies, arranged in a square grid, illuminated by two different light sources placed at N and E. This arrangement of light sources creates a gradient in intensity. Cells stochastically switch between biasing their motion towards one of the two light sources, with the effective intensity of the light sources towards N and E represented by *p*_1_ and *p*_2_ respectively. The map of *p*_1_ and *p*_2_ across a 5 × 5 grid is represented in (a) and (b) respectively. The morphology of colonies placed on this 5 × 5 grid is shown in (c). Each colony contains 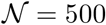 cells, and the model was simulated for 4 × 10^4^ time steps.

**Figure 5:**
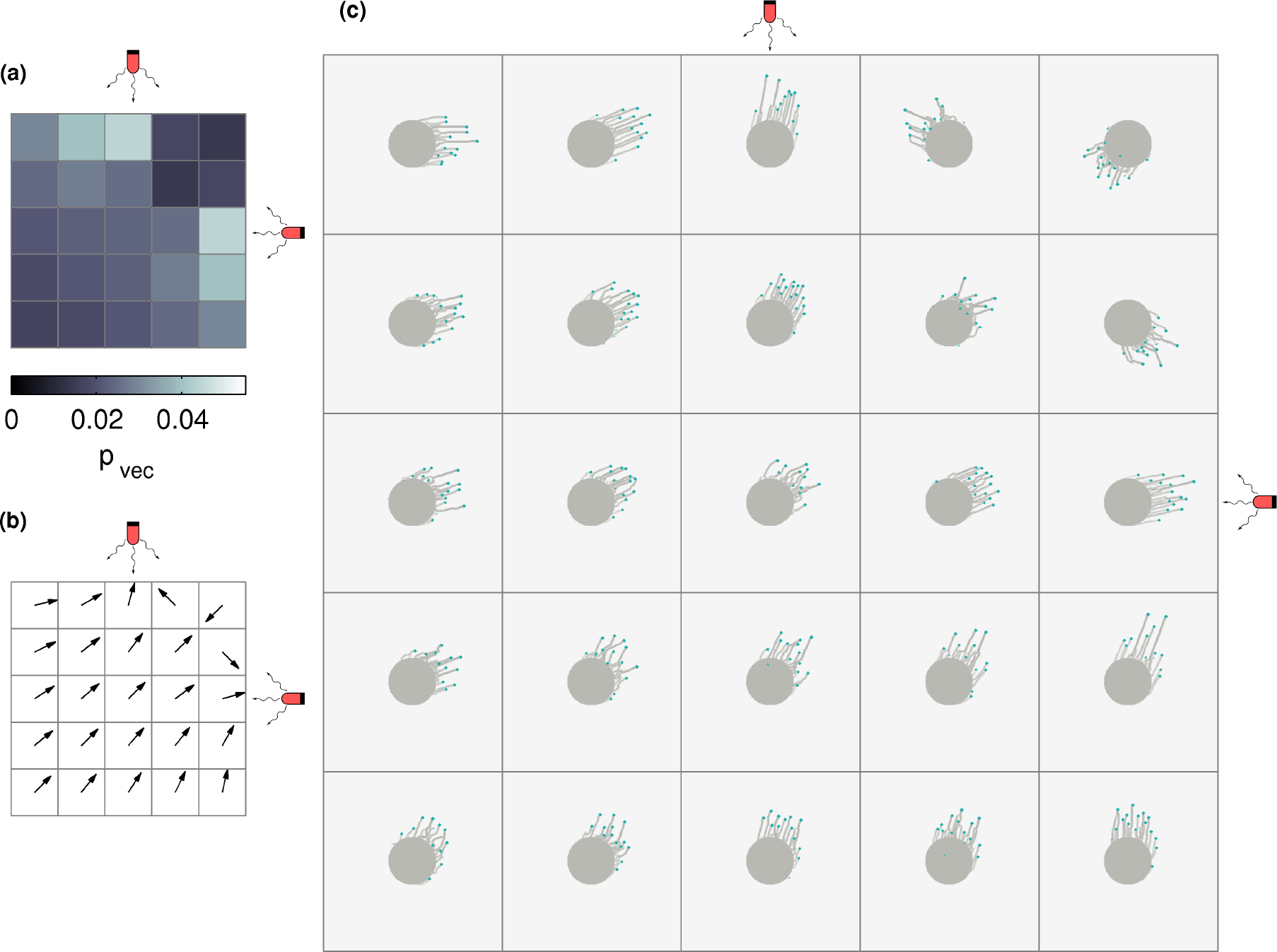
Cell colonies in a square grid under complex illumination in the vector integration case. Configurations of a number of cell colonies, arranged in a square grid, illuminated by two different light sources placed at N and E. This arrangement of light sources creates a gradient in intensity. Cells bias their motion towards the vector sum of the two light sources. (a) shows the effective total intensity of the light sources in the direction of the vector sum (*p*_vec_) is represented across a 5 × 5 grid. The direction of the vector sum of the light sources at each of the grid points is shown in (b) The morphology of colonies placed on this 5 × 5 grid is shown in (c). Each colony contains 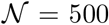 cells, and the model was simulated for 4 × 10^4^ time steps.

### Stochastic switching

The intensity of the light from the North and East experienced by a colony is captured by *p*_1_ and *p*_2_ respectively, using the convention of Fig. 1(b). Fig. 4(a) and Fig. 4(b) show the variation of light intensity (as expressed in terms of *p*_1_ and *p*_2_) over the colony array. Within each colony the *p*_1_ and *p*_2_ are constant. Cells sense these separate light sources and, at each time step, make a stochastic decision to move towards either light source, weighted by these probabilities. Each cell can thus decide to move towards North and East with probability *p*_1_ and *p*_2_ respectively, or can move in a random direction, chosen uniformly from [0, 2*π*], with probability 1 − *p*_1_ − *p*_2_.

As expected, and as shown in Fig. 4(c), colonies extend more pronounced fingers towards the closer light source. Colonies that are equidistant from each source extend fingers in the general N-E direction but appear to meander more. Note that the colony at the N-E corner of the grid extends fingers towards the S-W because the light sources are South and West of it.

### Vector integration

As in the case of stochastic switching, the intensity of the light experienced by a colony from the North and East is captured by the probabilities *p*_1_ and *p*_2_ respectively. Fig. 5(a) shows this map of probabilities *p*_vec_, experienced by each colony, where *p*_vec_ arises from the vector sum of *p*_1_ and *p*_2_. Fig. 5(b) shows the direction *θ*_vec_ of the vector sum. Within each colony the *p*_vec_ and *θ*_vec_ are constant. At each time step, cells make a decision to move in the direction of *θ*_vec_, with a probability of *p*_vec_ and can move in a random direction, chosen uniformly from [0, 2*π*] with probability 1 − *p*_vec_. As seen in Fig. 5(c), the qualitative nature of all colony morphologies are very similar to those for the stochastically switching case. The qualitative dependence of finger properties on intensity remain the same as in the one-dimensional case (see Fig. 3).

Comparing these two cases, we note that while the decision making process of individual cells is different, the final colony morphologies are strikingly similar. We reason as follows: although individual cells may decide to move according to one rule or the other, the overall morphology of the colony is a collective property arising also from the dynamic interaction between moving cells. This similarity suggests that observations of gross colony morphology may not suffice to disentangle the underlying mechanism of phototaxis at the single cell level.

To understand how individual cells integrate information in scenarios where they are exposed to complex illumination, we consider the information that can be extracted from individual cell trajectories.

### Distinguishing stochastic switching and vector integration scenarios

We considered a series of arrangements of light sources in order to systematically investigate the behaviour of cell trajectories under the different scenarios of stochastic switching and vector integration. We study a colony in which cells experience light from two sources that are placed on a circle centered at the colony. This ensures that the cells experience the same intensity of light, regardless of the angle that the light source makes with the East-West axis.

We consider three cases, where a pair of light sources are placed 30°, 60° or 90° North/South of East, respectively. In each case, we track the trajectories of individual cells over time, computing the angles that the trajectory makes, over each unit time interval, with the horizontal. In Fig. 6, we visualize the distribution of these angles as rose plots, where the height of each bar represents the relative probability that cells move in that angle.

**Figure 6:**
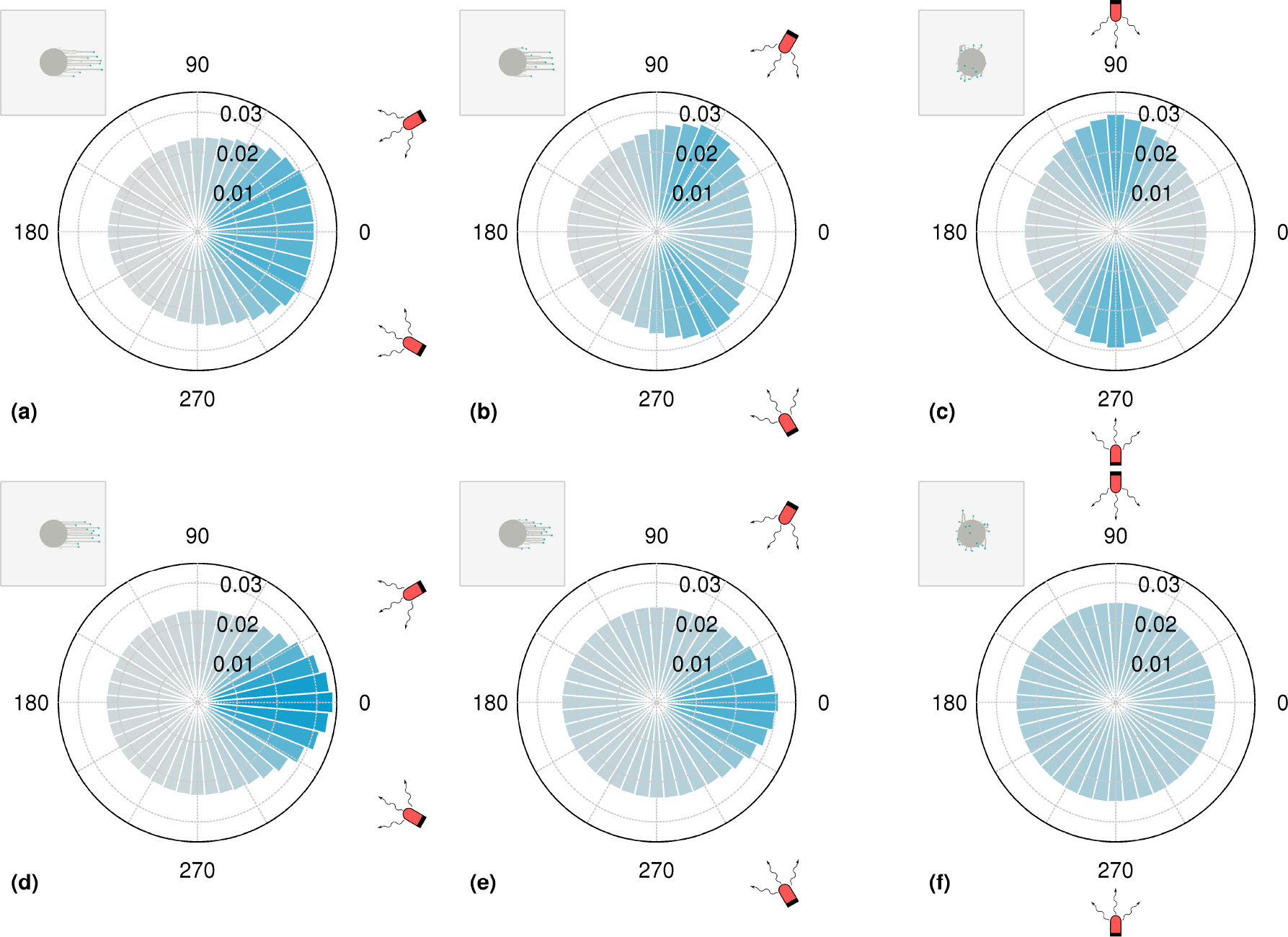
Rose plots of the angle of motion for stochastic switching and vector integration cases. Distribution of angles of motion of cells in colonies exposed to a pair of light sources placed at different positions relative to it. The probabilities are represented by the histogram of the angles made by the movement of a cell in a single time step. The light sources are placed at angles 30° (a,d), 60° (b,e) and 90° (c,f) North/South of East as shown in the respective figures. The intensity (and related *p*_photo_) is constant across all figures and is also assumed not to vary significantly across the size of the colony. In (a-c) at each step cells stochastically bias their motion in the direction of one of the light sources. In (d-f) at each step cells bias their motion towards the vector sum of the light sources. In all cases, the intensity of color of the bars of the rose plots are related to their magnitude. In each subfigure, the insets show the final colony morphology. Each colony contains 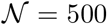 cells, and the model was simulated for 4 × 10^4^ time steps for the case *p*_1_ = *p*_2_ = 0.05.

Fig. 6(a-c) represents the stochastic switching case, where at each step cells stochastically bias their motion in the direction of one of the light sources. Fig. 6(d-f) shows results for the vector integration case, where at each step cells bias their motion in direction of the vector sum of the light sources.

In the case where the cells stochastically switch between detecting the two light sources, we find that the rose plots are characterized by two clear peaks in the directions of the light sources. This is in clear contrast to the corresponding rose plots obtained for the vector integration case, where cell motion is biased in the direction of the vector sum of the direction of the two light sources.

As the angle between the two sources is increased in the vector integration scenario, the cancellation of the effects due to the two opposing sources becomes more prominent. In contrast, the rose plots in the stochastic switching scenario indicate a clear bias in the direction of each individual light source. This difference between the two scenarios is particularly prominent in the case where the light sources are placed 90° North/South of East. Here, we observe two peaks that point in opposite directions for the stochastic switching case, while for the vector integration case we find that the distribution of angles in the rose plots is nearly uniform.

### Response of colonies with mixtures of cell types sensitive to different wavelengths

Cell colonies can be heterogeneous, expressing different levels and types of light receptors. The collective response in such communities can thus be influenced by the relative proportions of cells that respond differently to complex illumination.

In Fig. 7, we show results from our simulations for colonies consisting of varying proportions of cells that are sensitive to either red or green light. The simulated colonies are subjected to red and green light sources placed at different angles from the East-West axis. The ratios green-light sensitive cells to red-light sensitive cells are varied across 50:50, 25:75 and 10:90. Since each cell detects only one light source, the distinction between stochastic switching and vector-integration scenarios is inapplicable.

**Figure 7:**
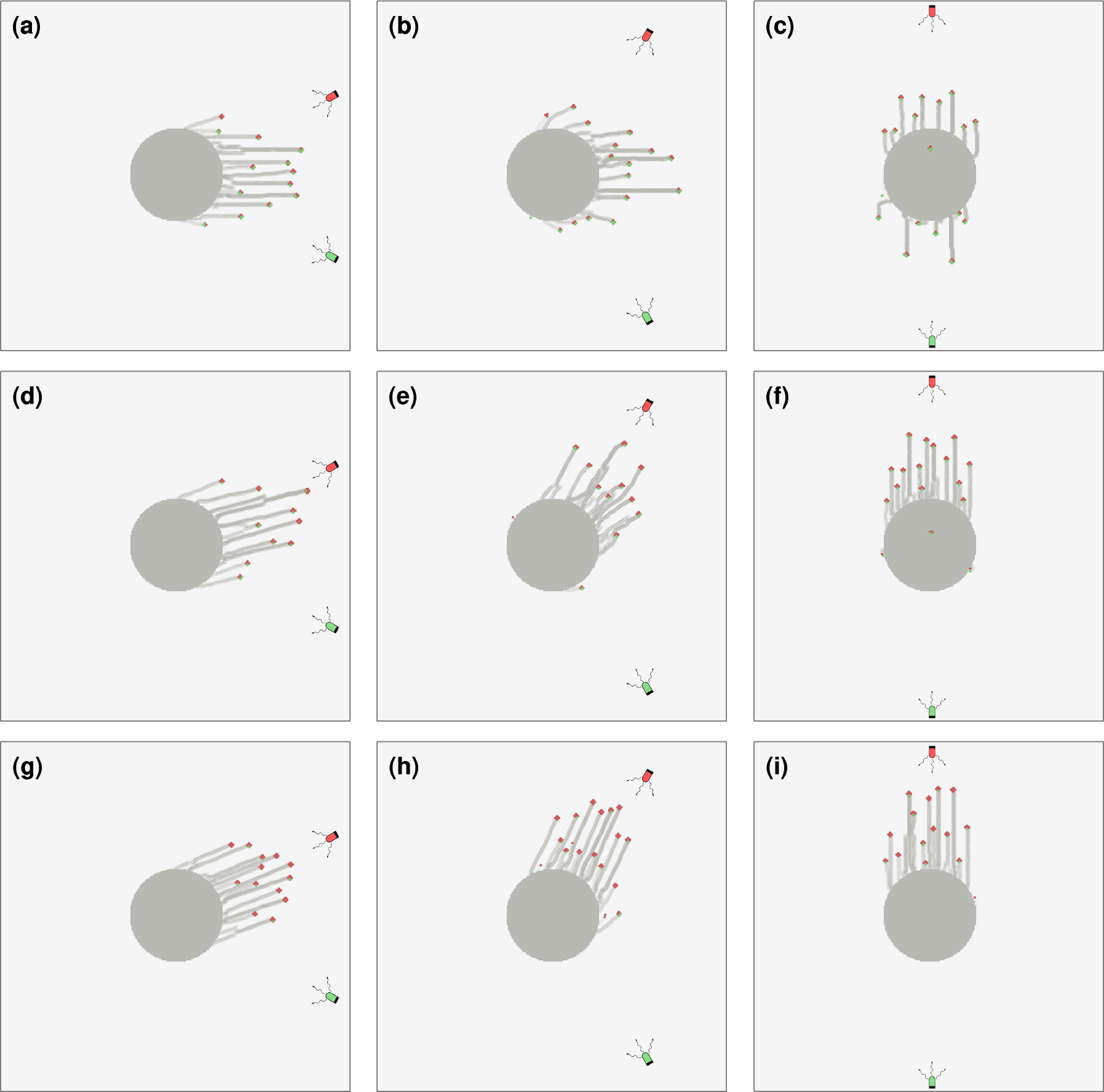
Colonies with mixtures of of cells sensitive to different wavelengths. Colonies are assumed to consist of two types of cells, each of which senses only one of the light sources. The cell types are indicated by green and red colors, corresponding to the wavelength of light they are sensitive to. The two different light sources, of the same intensity (and hence same *p*_photo_), are placed at angles 30° (a, d, g), 60° (b, e, h) and 90° (c, f, i) North/South of East as shown in the respective figures. We consider the following proportions of cells that sense only red to those that sense only green: (a-c) 50:50 (d-f) 75:25 (g-i) 90:10. Each colony contains 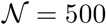 cells, and the model was simulated for 4 × 10^4^ time steps for the case where *p*_photo_ = 0.05 for both the red and green senstitive cells.

We start from an initial configuration where cells sensitive to different wavelengths are seeded at random. As the proportion of cells sensitive to red light is increased, the fingers are directed more towards the red light source. However, green-sensitive cells are also incorporated into these fingers as a result of cell-cell attachments. The green-light-sensitive cells in these fingers are not located at random, but tend to be found closer to the green light source. Finally, colonies consisting of roughly equal proportions of green and red-light sensing cells, tend to have more irregular fingers, an effect that is more prominent at intermediate angles between the two sources.

### Negative phototaxis

Finally, we study the response of cell colonies to light sources that induce a negative phototactic response in cells. Experimentally, illuminating colonies directionally under UV light has been shown to lead to the formation of fingers extending in a direction opposite to the light source [9]. We simulate a negative phototactic response through a probability that cells in the colony attempt to move away from the light source *k*, in the direction Θ_*k*_ − 180°, with the probability *p*_photo_. As before, there is also a random component to the direction of motion and the attachments of cells dictate collective colony morphologies.

In Fig. 8, we show results for negative phototaxis in initially circular cell colonies illuminated by UV light that is incident from the East. As in Fig. 3, the length and rate of growth of fingers depends strongly on *p*_photo_, with the difference that the fingers now extend away from the source rather than towards it.

**Figure 8:**
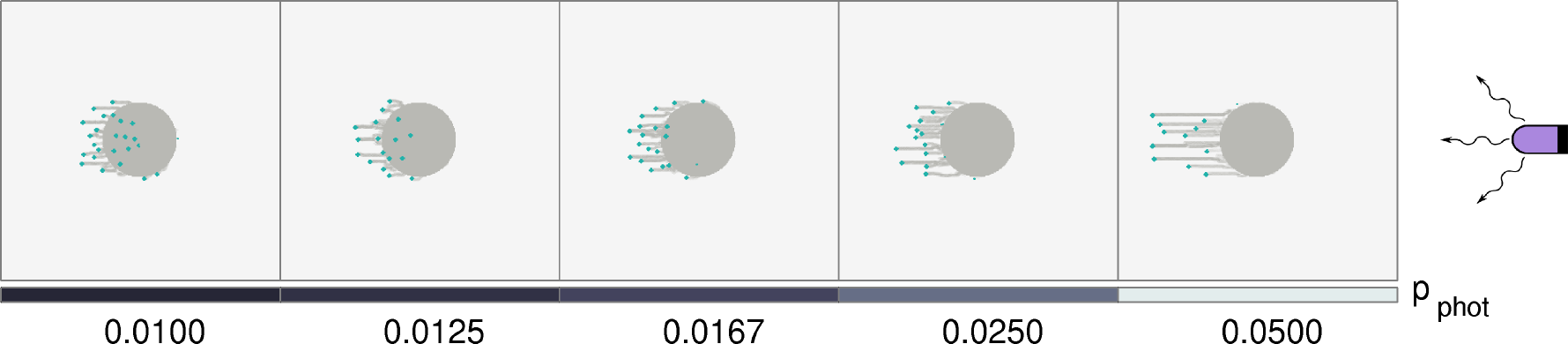
Colony morphology with negative phototaxis (UV light). The colony farthest to the East is the closest to a UV light source, and experiences the greatest light intensity, and hence the largest *p*_photo_. As shown in the bottom panel, *p*_photo_ decreases linearly towards the West. At each time point, an individual cell can either detect the UV light and move away from it, or can move in a random direction. Each colony contains 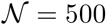 cells, and the model was simulated for 4 × 10^4^ time steps.

## Discussion

In this paper, we explored the consequences of complex illumination on the shapes of cyanobacterial colonies. We used a computational model whose fundamental unit was the single cell. We modeled each such cell as an agent whose movement was dictated by its own interactions with external light sources, as well as from its interactions with its neighbours. The motion of the agent was facilitated by the slime that it encountered when moving. The model accounts explicitly for the forces exerted, and experienced, by each cell due to its neighbours through attachments mediated via T4P.

We studied the morphologies of colonies under different light regimes to determine the mechanisms through which single cells integrate information from external cues, translating these into decisions regarding their motion. We investigated at least two major mechanisms by which phototactic cells could respond to light incident on them from different sources. The first was a stochastic switching scenario in which cells chose, at each time step, either to move towards a randomly chosen light source with a fixed probability, or to move in an arbitrarily chosen direction. In the second model, cells responded through what we term “vector integration”, in which individual cells either chose to move along the vector sum of individual light sources, or chose, at each time step, to move in a random direction. The specific scenario applicable could also, in principle, depend on the wavelengths of light as well as the magnitude of the intensity that the cell is exposed to. We asked if the subtle difference between these scenarios, clearly distinct at the single cell level, could be inferred from large-scale measurements on colonies.

We observed that similar-looking colony morphologies could be obtained under two very different underlying mechanisms of single-cell response to complex illumination. To explore this further, we examined the statistics of the trajectories of individual cells within the colony. We concluded that extracting statistical features of individual cell trajectories could provide a way of distinguishing between these, and potentially other, mechanisms of single-cell response, even if the overall colony morphologies did not. We describe ways in which the relevant information can be extracted from ensembles of single cell trajectories.

At a more general level, colonies of phototactic bacteria provide a unique opportunity to test models of collective response in living systems. While individual bacteria can sense and move towards light [10], the nature of colony morphologies is a function of the mechanical attachments between cells, mediated by their T4P [25], as well as of the slime that cells lay down [26].

Slime, in particular, plays a unique role. It allows for density-dependent motility [27], reminiscent of quorum-sensing mediated by small molecules in bacteria. Unlike quorum-sensing, however, slime-mediated interactions between cells can be time-dependent, since the motion of bacteria at later times can be influenced by slime laid down by other bacteria at an earlier time. In conventional quorum-sensing, driven by the production and detection of small diffusible autoinducers [28], the relatively large diffusion constant of small molecules implies that the time delays between production and detection can be safely ignored.

Swimming bacteria such as *E. coli* interact via a practically instantaneous and long-ranged hydrodynamic interaction mediated by the fluid [29, 30]. This is a feature of virtually all biophysical models for the interactions between swimming bacteria. In contrast, the interactions between cyanobacteria are short-ranged, involve direct mechanical forces mediated by T4P, and could also be delayed in time. Thus, phototaxis differs in a number of qualitative ways from the much-studied problem of bacterial chemotaxis.

Computational models such as the one we describe here, once benchmarked, can be used to investigate behaviour that can be experimentally tested in order to identify regimes of the parameter space that are most likely to provide strong evidence for one hypothesis over another. Such models can also be used to study behaviour in other types of bacteria that interact with each other using T4P. The agent-based approach discussed here makes it easy to incorporate inter-cellular variations in response to stimuli, such as in colonies comprising mixtures of cells that respond only to a particular incident wavelength. These may also naturally correspond to the situation in heterogeneous ecological communities of phototactic bacteria, such as in hot springs [31].

Additional detail can be incorporated into the model we describe here, allowing us to bridge the gap between single cell and collective response more effectively, as more data becomes available. Finally, there are fascinating, and as yet unanswered, questions that relate to how cells in bacterial colonies might localize pili and photoreceptors to activate downstream signaling pathways. Coupling such signaling responses to cell-cell interactions and cell motion would enable us to address a number of questions concerning the mechanobiology of collective cell behaviour.

## Author Contributions

SNM and PV carried out simulations. SNM, PV and GM designed the project and prepared the manuscript.

## Acknowledgements

SNM is supported by the IMSc Complex Systems Project (12^th^ Plan). We would like to thank Devaki Bhaya for helpful discussions and comments on the manuscript.

