## Supplementary Material for "Information Integration and Collective Motility in Phototactic Cyanobacteria"

#### Caption for Movie

**Movie S1**: Time lapse movie showing the positions of cells and the slime they lay down over the course of a simulation. We start with an array of colonies under a single light source placed to the East of the colonies as described in Fig. 3. Here $p_{photo}$ decreases linearly towards the West, as shown in the bottom bar. Each subsequent frame is separated by 500 time steps. Each colony contains $N = 500$ cells, and the model was simulated for $4 \times 10^4$ time steps.

### Figures

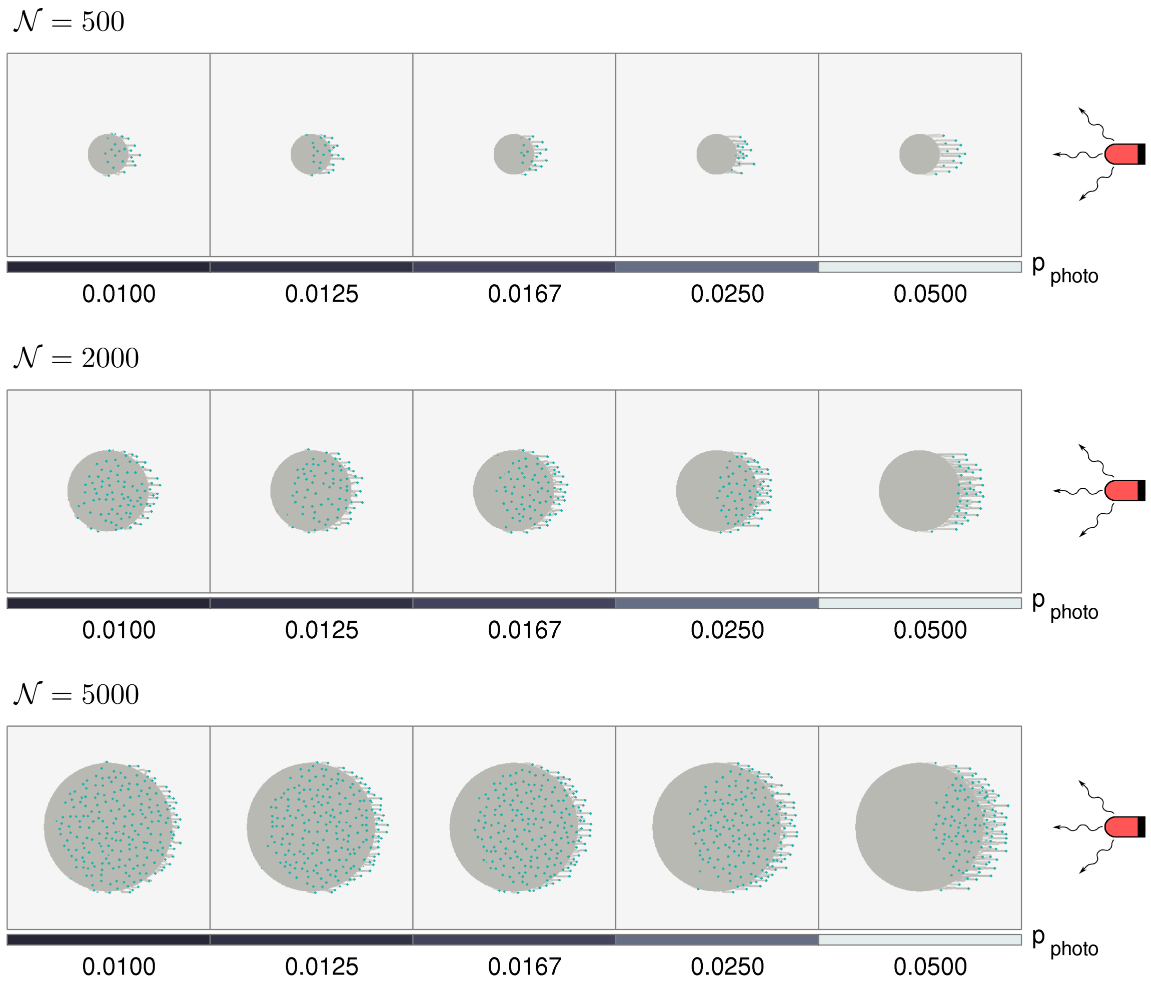


Figure 1: **Morphologies of colonies in a linear grid under structured illumination obtained for different values of $\mathcal{N}$.** Colonies are illuminated by a single light source placed at the East. Each experiences a different intensity of incident light, depending on its location relative to the light source, such that the easternmost colony experiences the greatest incident light intensity. Thus, $p_{\text{photo}}$ decreases linearly towards the West, as shown in the bar below each row. The number of cells in each row is indicated by $\mathcal{N}$, and the model was simulated for $3 \times 10^4$ time steps. Note that in each of the panels we have used a slime matrix of dimension $400 \times 400$, which is larger than those used for figures in the main text ($300 \times 300$).

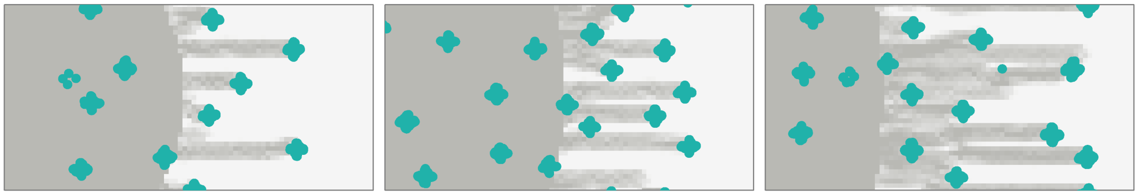

$\mathcal{N} = 500$ $\mathcal{N} = 2000$ $\mathcal{N} = 5000$

Figure 2: **Morphologies of colonies obtained for different values of $\mathcal{N}$.** Morphologies of three colonies, each under illumination from a single light source placed at the East. The number of cells in each colony is (left-right) $\mathcal{N} = 500$, $\mathcal{N} = 2000$ and $\mathcal{N} = 5000$ cells, as indicated below the corresponding panels. Each colony experiences a light intensity of $p_{\text{photo}} = 0.05$ and the model was simulated for $3 \times 10^4$ time steps. In each case, we display a close-up of the morphologies around the colony edge, where the dimensions of each box is identical. The displayed results indicate that the morphology of the fingers is relatively independent of the number of cells $\mathcal{N}$.

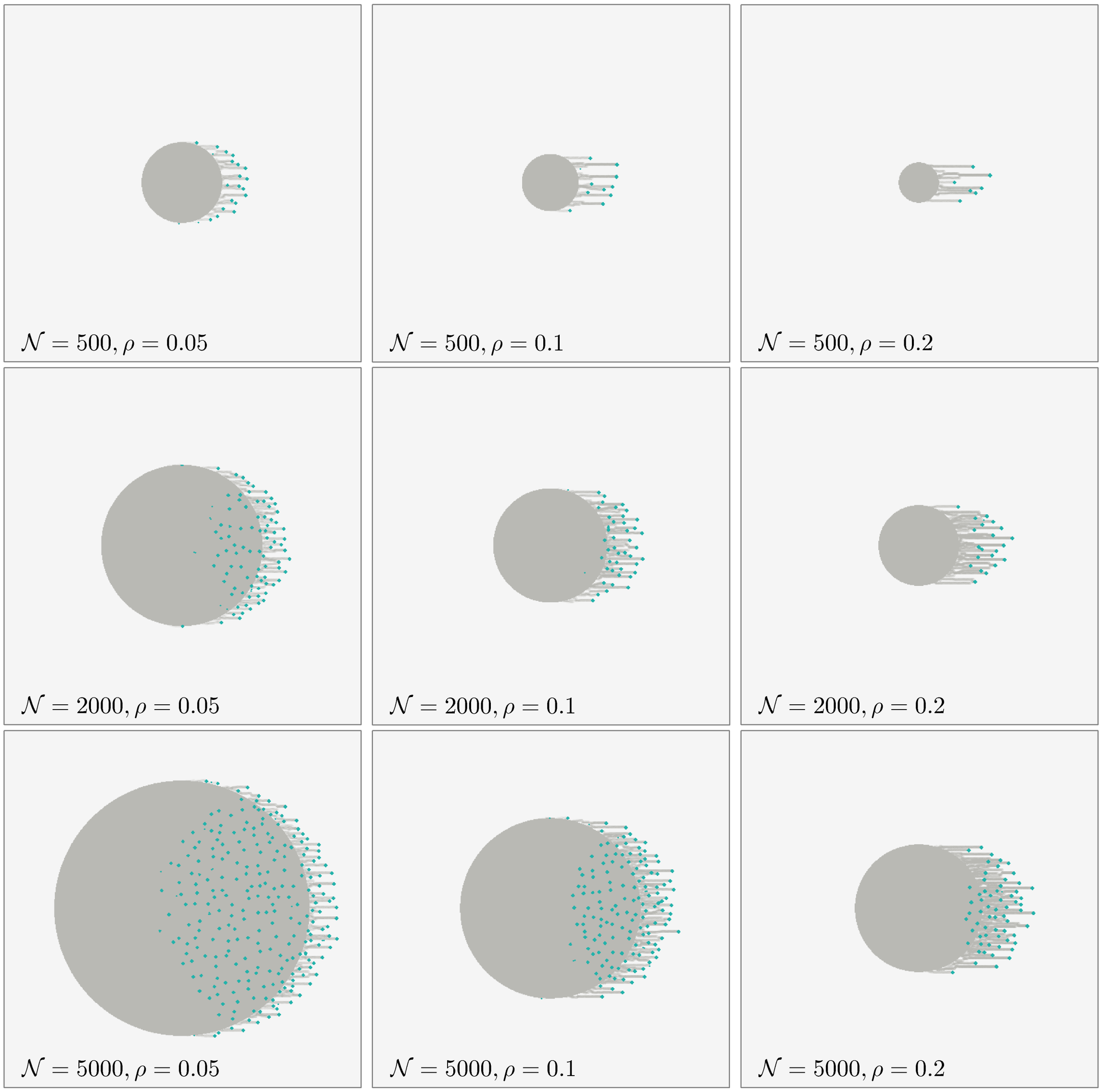


Figure 3: **Morphologies of cell colonies of different values of $\mathcal{N}$ and densities $\rho$.** Morphologies of several colonies under illumination from a single light source placed at the East. Each colony experiences a light intensity of $p_{\text{photo}} = 0.05$, the number of cells in each panel is indicated by $\mathcal{N}$, the colony density is indicated by $\rho$ and the model was simulated for $3 \times 10^4$ time steps. Note that in each of the panels we have used a slime matrix of dimension $500 \times 500$, which is larger than those used for figures in the main text ($300 \times 300$).
