## Supplementary figures and images for "Information Integration and Collective Motility in Phototactic Cyanobacteria"

### Movie S1

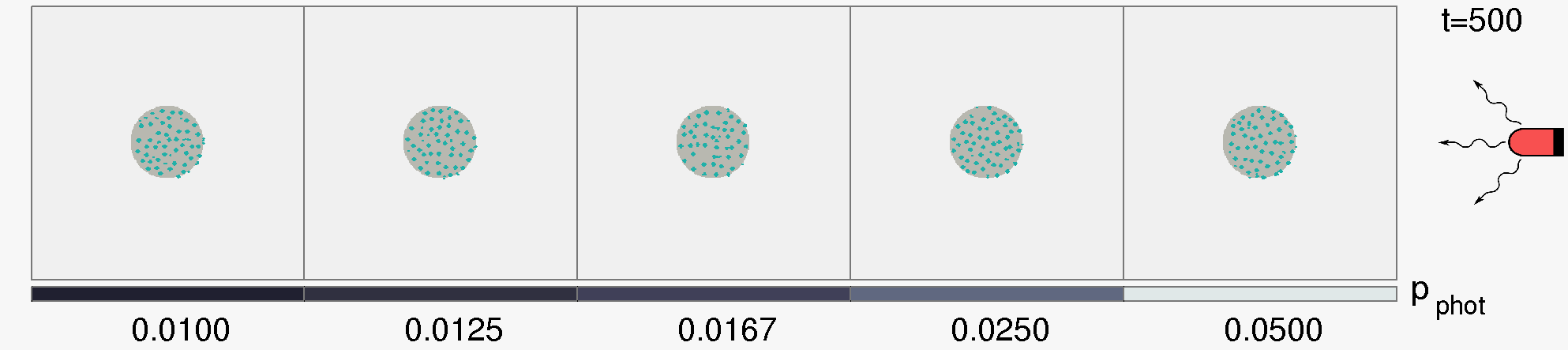
